# Unravelling the role of a disordered chaperone in adaptation to environmental stress

**DOI:** 10.64898/2026.06.01.729257

**Authors:** Deepak T Hurali, Anand Ballal, Manisha Banerjee

**Affiliations:** Molecular Biology Division, Bhabha Atomic Research Centre, Mumbai 400085 India; Nuclear Agriculture & Biotechnology Division, Bhabha Atomic Research Centre, Mumbai 400085 India; Homi Bhabha National Institute, Anushakti Nagar, Mumbai 400094, India

## Abstract

Intrinsically disordered proteins (IDPs) lack a stable tertiary structure, which enables them to mediate flexible molecular interactions. As the biochemical functions of IDPs remain poorly understood, their physiological roles are largely unknown, particularly in photosynthetic organisms. Herein, Alr0806, a conserved salinity-induced cyanobacterial IDP, was characterized from the nitrogen-fixing cyanobacterium *Anabaena*. Instead of the full-length protein predicted in databases, experimental analysis indicated this organism to express a shorter form of the Alr0806 protein, which was attributed to the mis-annotation of the translational start codon. Purified Alr0806, a highly thermostable protein, exhibited characteristic properties of highly disordered proteins, including anomalous migration on SDS-PAGE. Alr0806 displayed disorder-to-order transitions with decreasing pH, indicating structural flexibility. Notably, Alr0806 leveraged structural plasticity to function as a chaperone and molecular shield, protecting proteins from aggregation. Furthermore, consistent with this function, *Anabaena* strains deficient in Alr0806 showed compromised growth and diminished photosynthesis under standard conditions of growth or in response to salt/heat stress. These findings establish Alr0806 as a key player in cyanobacterial physiology and provide insights into the physiological functions of IDPs in photosynthetic organisms.

**Highlight:** Alr0806, a conserved cyanobacterial stress-induced intrinsically disordered protein, exhibits structural plasticity and functions as a molecular chaperone to support growth, photosynthesis and environmental stress tolerance.

## Introduction

Photosynthetic organisms continually face environmental perturbations (temperature extremes, water/nutrient deficits, ioinic challenges etc.) that negatively influence growth, development, and productivity. Such adverse conditions result in unfolding, denaturation, aggregation or degradation of proteins, ultimately compromising cellular fitness. Cells synthesize diverse classes of proteins that include enzymes, structural proteins, chaperones etc. to counter these stresses. More recently, the catalogue of stress-responsive players has expanded to include unstructured or intrinsically disordered proteins (IDPs), which consist of disordered regions and lack a well-defined tertiary structure. Late embryogenesis-abundant proteins (LEA) and dehydrins are some of the key stress-responsive IDPs characterized in plant systems (**Hsiao et al., 2024).**

IDPs can adopt different conformations, enabling their interaction with various partners. These proteins leverage their disorder-to-order transitions to bind membranes, enzymes, DNA, and metal ions, facilitating resilience to various stresses. Some IDPs protect cellular proteins by preventing their aggregation, particularly under stressful conditions. High net-charge, low hydrophobicity, low sequence complexity are the important characteristics of IDPs. As the function of IDPs depends heavily on electrostatic interactions, properties such as their isoelectric point (pI) assume considerable importance for their physiological roles (**Mitic et al., 2018**).

Cyanobacteria, the progenitors of plant chloroplasts, are widely used as model systems for plants as they possess conserved pathways related to photosynthesis, carbon fixation, and stress adaptation. Moreover, their adaptive responses to diverse abiotic stresses such as heat, salinity, dehydration, high light, and oxidative stress, resemble those observed in plants. The nitrogen-fixing cyanobacterium *Anabaena* PCC 7120 known for its ability to withstand multiple environmental stresses by inducing the synthesis of protective compounds, chaperones, and antioxidant systems, which mitigate stress-induced cellular damage (**Banerjee et al., 2012, 2013, 2015, Chakravarty et al., 2016, 2020**). In addition, *Anabaena* spp. holds considerable agricultural and economic importance because of their role as natural biofertilizers in rice fields.

In general, the study of IDPs especially regarding their *in vivo* role in stress adaptation is still in its nascent state. Notably, in *Anabaena* PCC 7120, many IDP encoding genes are upregulated in response to salt stress, indicating their importance in overcoming salinity. RNAseq data revealed the *alr0806* gene, which encodes unknown/uncharacterized IDP, to be upregulated under salt stress (**Hurali et al., 2024**). Induction of Alr0806 in response to heavy metals, drought and other stresses (**Srivastava et al, 2023; Agrawal et al., 2014; Sen et al., 2017**) lends credence to the hypothesis that this protein constitutes a crucial component of the cyanobacterial stress-adaptive machinery. But, despite its obvious importance, Alr0806 has not been physiologically or structurally characterized so far. This study, for the first time, has attributed a function to Alr0806, demonstrating that its works as a chaperone. CRISPRi-based knockdown analysis showed the Alr0806 to be essential for normal cellular growth as well as adaptation to environmental stress.

## Results

### 1. The actual ORF of the monocistronic stress-inducible gene *alr0806* is shorter than the ORF annotated in databases

The *alr0806* gene was strongly induced by salts (NaCl and NH Cl), but not by oxidative stressors (methyl viologen/H_2_O_2_) (**Figure 1A and Supp Fig.1A).** Northern blotting-hybridization analysis (**Figure 1B**) showed distinct induction of ∼0.5-kb transcript in filaments treated with NaCl or NH_4_Cl, indicating that the *alr0806* gene was monocistronic. Withdrawal of salt stress led to rapid disappearance of the *alr0806* transcript, demonstrating the requirement of this stressor for its continued transcription. The RNA seq data of *Anabaena* PCC 7120 (**Hurali et al., 2024**) indicated the probable transcriptional start site (TSS) of *alr0806* to be in between the (rare) translational start codon ‘GTG’ annotated in databases and the predicted translational start codon “ATG” (**Figure 1C**). When cDNA-based PCR assays were performed employing two distinct forward primers, corresponding to annotated (P1) or predicted translational start site (P2) and a common reverse primer (P3), amplification was observed only with the downstream primer-reverse primer pair (P2–P3) (**Figure 1C, inset)**. This finding implied that the actual translational start codon was likely to be ‘ATG’ that was 66 nucleotides downstream of the annotated ‘GTG’ start codon (**Figure 1C).** Thus, with the ATG start codon, the amino acid length of Alr0806 reduced from 157 to 135 amino acids.

**Figure 1.**
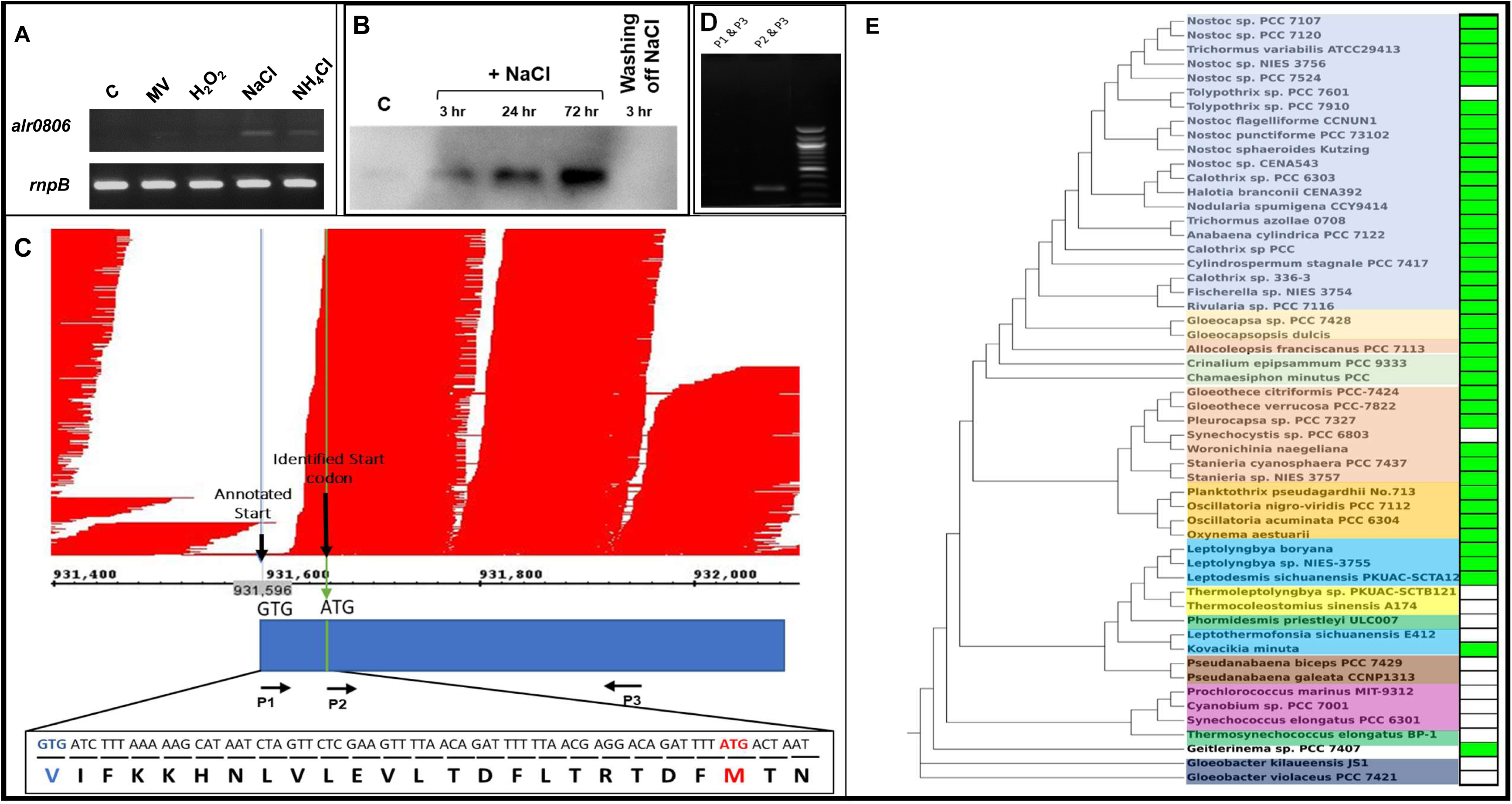
(A) Semi-quantitative RT–PCR analysis of *alr0806* under different conditions. *RnpB* served as the internal control. (B) Northern blot analysis. Total RNA from *Anabaena* cells treated with 100 mM NaCl was isolated at different time points (3, 24, and 72 h). After 72h, cells were washed, inoculated in medium without NaCl, and RNA was extracted after 3 h. After resolving the samples on denaturing agarose gels, RNA was transferred onto a nylon membrane and hybridized to the DIG-labeled *alr0806* probe. (C) Alignment of paired-end transcript reads onto the *Anabaena* PCC 7120 genome (GCF_000009705.1), visualized using Integrated Genome Browser. Red peaks indicate transcript coverage. The annotated translational start site, transcriptional start site, and newly identified translational start site is denoted. Horizontal arrows (P1, P2, and P3) indicate the positions of primers used for PCR. Inset: PCR amplification using cDNA as template with primer combinations P1–P3 and P2–P3. (D) Distribution of Alr0806 homologs across 54 cyanobacterial species. Green box indicates the presence, while white box indicates the absence of the Alr0806 homolog. The phylogenetic tree was constructed based on 16S rRNA sequences of the selected cyanobacteria.

Although the Alr0806 protein is denoted as an unknown protein in the NCBI database, it was referred to as ‘high light inducible protein (HLIP)’ homolog in earlier studies (**10**). Homology searches revealed very low overall homology with the other HLIPs, but a small stretch (93 to 110 amino acids) showed high similarity (66.67%) with the corresponding residues from *Synechococcus* sp. JA-3-3Ab HLIP (**Suppl Fig 1C**). Alr0806 homologs were predominantly present in morphologically complex, filamentous cyanobacteria, particularly among the heterocystous order of Nostocales (*Nostoc*, *Trichormus*, *Anabaena* etc.). Although these homologs were also recorded in several non-heterocystous filamentous taxa (e.g. *Oscillatoria* & *Leptolyngbya*), no Alr0806-like proteins were detected in the primitive cyanobacterial genus *Gloeobacter* or unicellular, genome-rationalized cyanobacteria, such as *Synechococcus* and *Synechocystis*. The presence of Alr0806 homologs broadly aligned with the major phylogenetic clades, but interestingly, several closely related taxa showed divergence, indicating that Alr0806 evolved later during cyanobacterial expansion and underwent lineage-specific retention or loss (**Figure 1D).**

### 2. The purified Alr0806 protein exhibits typical characteristics of disordered proteins

Interestingly, disorder tools predicted the database-annotated Alr0806 to have ∼62% overall disorder (**Figure 2A).** The first 27 amino acids and an amino acid stretch from 83-116 residues was ordered (**Figure 2A**), whereas the rest of the protein (28-82 and 117-157 amino-acid stretches) was disordered. However, in the 135 amino acid Alr0806 protein, due to the absence of the ordered 27 amino acid stretch, the overall disorder increased to 76%. AlphaFold predictions of Alr0806 showed the presence of ordered as well as disordered regions (**Figure 2A, lower panel)**. This shorter 135 amino acid version of Alr0806, which is characterized in the subsequent sections, is henceforth referred to as the Alr0806 protein. This protein had a low isoelectric point (pI: 4.04), a high overall negative charge (-17.5) and inherent propensity to undergo liquid-liquid phase separation (LLPS) (**Suppl Figure 2A**). *In silico* physicochemical analysis categorized this protein in the “strong polyampholyte” category (**Supp Fig 2B**). Consistent with this analysis, the charge–hydropathy (CH) plot positioned Alr0806 firmly within the IDP region (**Figure 2B).**

**Figure 2:**
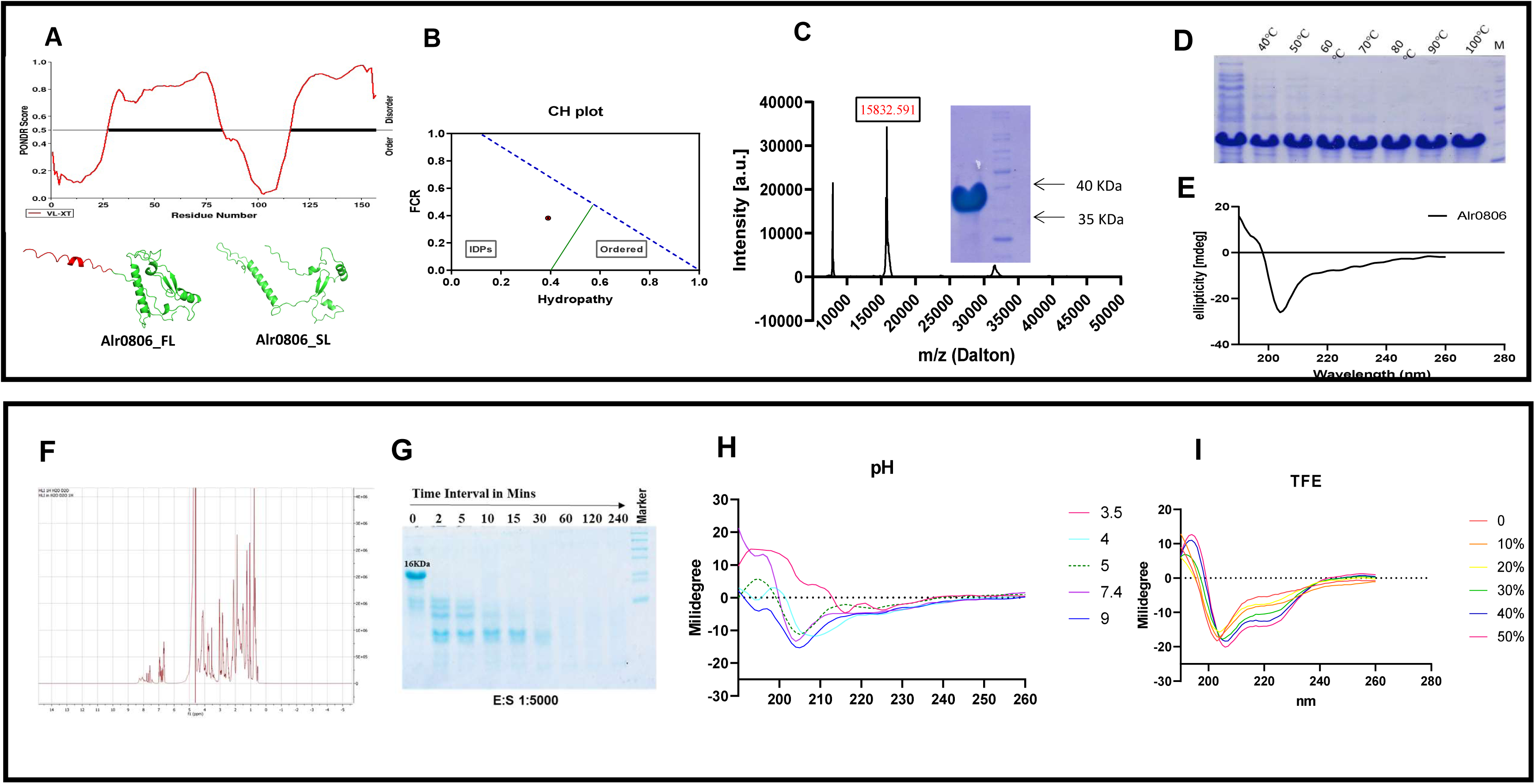
(A) Disordered prediction using PONDR for the 157 residue Alr0806 protein. Amino acid stretches with PONDR score ≥ 0.5 were considered as disordered. The lower panel shows the predicted Alpha fold 3D structure of both the 157 and 135 amino acid residue Alr0806. Red color indicates the misannotated region. (B) Charge-Hydropathy (C-H) Plot. This graph plots the mean net charge (FCR) against the normalized mean hydropathy for the Alr0806 protein. The standard boundary line (green line) separates the compact proteins (typically folded) from the more extended IDPs. (C) MALDI TOF Mass spectrometric profile of the Alr0806 protein. The SDS profile of the 135 amino acid residue Alr0806is shown in the inset. (D) Soluble fractions of cell lysates were subjected to different temperatures (as indicated) and analyzed on denaturing SDS–PAGE. M, molecular weight marker. (E) CD spectrum of Alr0806 (F) 1D NMR spectrum of Alr0806. (G) Limited proteolysis of Alr0806 by Proteinase K at E:S molar ratio 1:5000 st different intervals of time. (H) CD spectrum spectra of Alr0806 protein at different pH. (I) CD spectra of Alr0806 treated with different concentration of TFE (0-50% v/v).

The Alr0806 protein was overexpressed in *E. coli* with C-terminal 6 His tag and purified to near-homogeneity (**Supp Figure 2C**). The purified protein displayed anomalous migration on SDS–PAGE, appearing at considerably higher apparent mass of ∼35 kDa, instead of the expected 15.8 kDa (**Figure 2C, inset).** However, MALDI–TOF mass spectrometry authenticated the molecular weight of this protein to be ∼15.8 kDa, which exactly matched with its predicted molecular weight (**Figure 2C).** Even after exposing the cell lysate containing the overexpressed protein to 100 □C, Alr0806 remained soluble, demonstrating high degree of thermostability (**Figure 2D).** CD spectroscopy showed a strong negative ellipticity near 204 nm, which is typical for “random coil” or “disordered” proteins. At increasing temperatures too, no significant spectral changes, were observed, indicating exceptional thermostability and preservation of its disordered conformation (**Figure 2E and Suppl Fig 2D).**

To rule out the effect of the added his-tag on the protein’s structural properties, untagged version of Alr0806 was also overexpressed in *E. coli* and purified by subjecting the lysates to boiling followed by anion-exchange chromatography **(Supp Fig 2E)**. The untagged Alr0806 too showed anomalous migration on SDS-PAGE, but MALDI–TOF analysis ascertained its mol. wt. to be 15 kDa, which correlated with its expected molecular mass (**Supp. Fig. 2E inset**). Native PAGE analysis showed a single band, while dynamic light scattering (DLS) measurements too revealed a single monodisperse peak **(Supp Figure 2F and 2G**). Thus, both His-tagged and untagged Alr0806 proteins exhibited comparable biophysical characteristics. One-dimensional NMR spectra of Alr0806 revealed limited dispersion in chemical shift, which is consistent with a predominantly disordered conformation (**Figure 2F**).

### 3. The Alr0806 protein is very susceptible to proteolytic cleavage and shows disorder to order transitions

Limited proteolysis with Proteinase K showed rapid degradation of Alr0806, with only a few protease-resistant fragments persisting after 4 h at a low enzyme-to-substrate ratio (1:10,000) (**Figure 2G)**. These stable fragments are likely to correspond to structured domains within an otherwise flexible polypeptide, in agreement with *in silico* predictions of mixed ordered and disordered regions. CD analysis across different pH revealed an increase in helical content at pH 4 as compared to pH 7.4 (**Figure 2H**), indicating considerable alteration in conformation near its pI. The intrinsic tyrosine fluorescence data also showed a reduction in the peak fluorescence with varying pH, implying conformational changes at different pH **(Supp Figure 2H).** Notably, the addition of trifluoroethanol (TFE), a known helix-stabilizing agent, caused a significant increase in helicity at concentrations 30% (v/v) and above, denoting the presence of latent helical regions within the disordered protein (**Figure 2I).**

### 4. Alr0806 functions as a chaperone and knocking down *alr0806* decreases *in vivo* tolerance to salinity or heat

As the structural flexibility of Alr0806 is likely to allow it to interact with multiple proteins, the ability of Alr0608 to function as a chaperone was evaluated. In the concentration-dependent manner, Alr0806 effectively protected MDH from aggregation in thermal denaturation assays, demonstrating its holdase activity (**Figure 3A).** Similarly, MDH protein which is sensitive to dehydration, retained a higher proportion of enzymatic activity in the presence of Alr0806, suggesting that Alr0806 also functions as a molecular shield under water-depleted conditions (**Figure 3B).**

**Figure 3:**
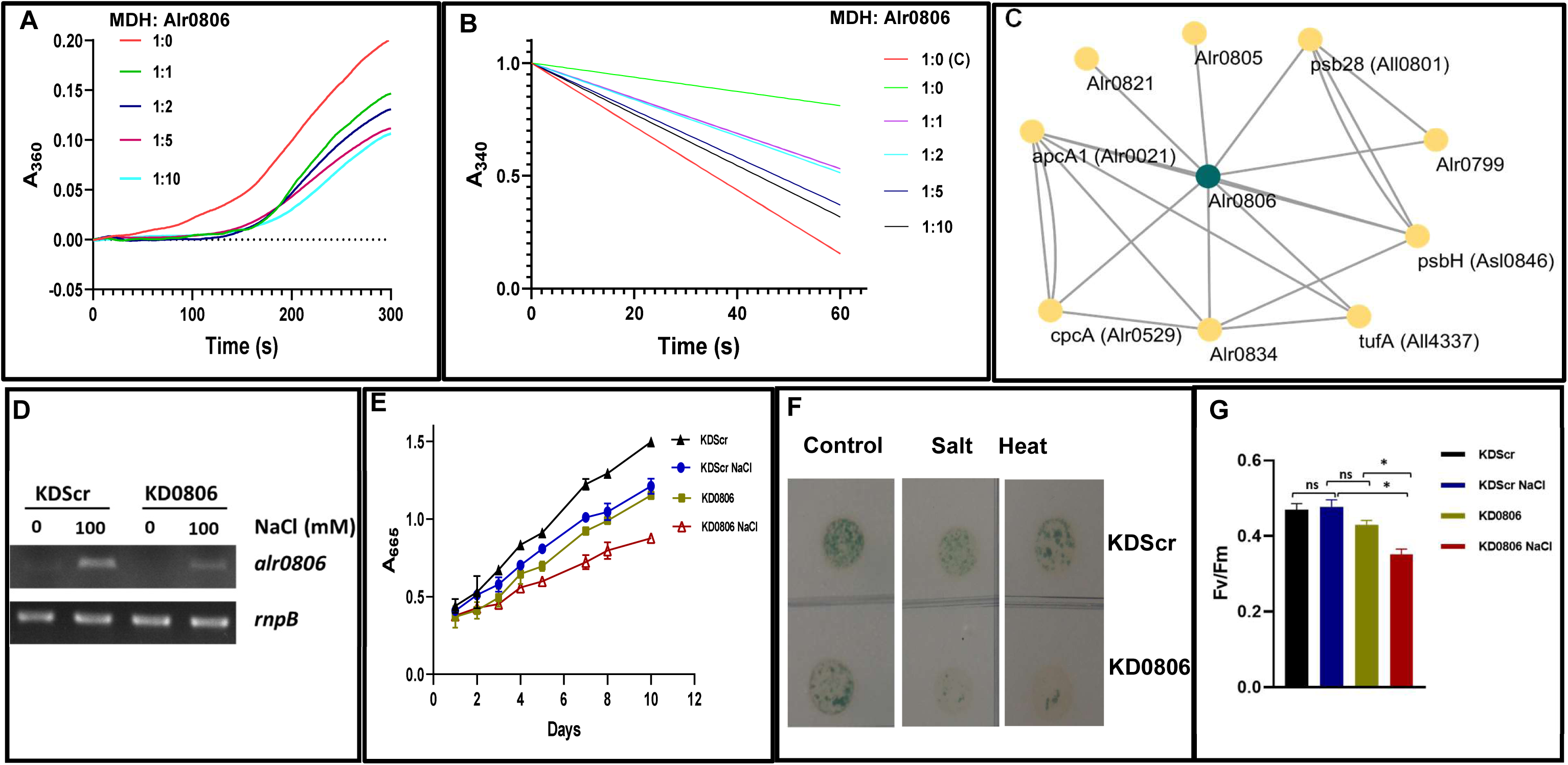
(A) Holdase activity of Alr0806 was assessed by monitoring the aggregation kinetics of malate dehydrogenase (MDH) at 55 °C under varying molar ratios of MDH to Alr0806(1:0, 1:1, 1:2, 1:5, and 1:10). (B) Dehydration protection assay showing the kinetics of NADH consumption by MDH after rehydration in the presence of different molar ratios of MDH to Alr0806 (C – control) (C) The PPI network for Alr0806 constructed using the Cytoscape tool (https://cytoscape.org). Green dot represents Alr0806 while the yellow dots represent the interacting proteins. (D) Quantitative reverse transcriptase PCR to monitor the content of the *alr0806* transcript in the control (An KDScr) or the knockdown strain (An KD0806). (E) Growth of An KDScr or An KD0860 with or without 100 mM NaCl. (F) On day 4 of the growth curve, An-KDScr and An-KD0806 aliquots were spotted on BG-11 plates and photographed after 12 days of growth under standard conditions. For heat stress, the cultures were subjected to 42°C (1 h) and subsequently spotted on BG-11 plates.

The protein–protein interaction (PPI) network of Alr0806 was constructed using the experimentally validated database CyanoMapDB (**Peng et al., 2023**). This analysis revealed Alr0806 to interact with 12 partner proteins (**Figure 3C)** involved in photosynthesis, thylakoid membrane organization, phycobilisome assembly, and other intracellular processes (**Supp. Figure 3A**). Furthermore, PPI analysis by **Xu et al. (2022)** had earlier demonstrated Alr0806 to form a larger complex comprising of 51 proteins (**Supp. Figure 3B**). Probably, the chaperone activity of Alr0806 aids in maintaining appropriate conformation and function of these interacting partners.

To assess the *in vivo* role of *alr0806*, a CRISPRi-based construct (pKD0806) was employed to knock down expression of Alr0806 in *Anabaena*. A corresponding construct featuring a scrambled sgRNA (KDScr) served as the control. After transferring these constructs into *Anabaena* PCC 7120 via conjugation, the knockdown nature of AnKD0806 was confirmed by RT PCR (**Figure 3D**). AnKD0806 demonstrated reduced growth and decreased photosynthetic parameters such as photosynthetic efficiency (F_V_/F_M_) in comparison to AnKDScr under standard conditions of growth or on treatment with NaCl (**Figure 3E**). Very likely, the association of Alr0806 with phycobilisomes or PSII complex constituent proteins (e.g. CpcA, PsbH & Psb23) is important for optimal photosynthetic efficiency. At day 4, when these cultures were spotted on BG-11 agar plates and incubated further, the knockdown strain showed decreased survival relative to KDScr, indicating heightened sensitivity to salinity (**Figure 3F**). It should be noted that even in the absence of salinity, AnKDScr showed better growth than AnKD0806, which correlates well with the growth analysis shown in Figure. 3E. Similarly, when the exponentially growing 4-day old filaments were subjected to heat shock and spotted onto BG-11 plates, the AnKD0806 strain showed diminished survival (**Figure 3F**). These findings highlight the critical role of *alr0806* in sustaining cellular homeostasis under salt or heat stress.

## Discussion

Intrinsically disordered proteins (IDPs) represent a fascinating yet relatively uncharted area of research within the discipline of molecular and structural biology (**Uversky et al., 2015)**. Despite their prevalence, the physiological roles of IDPs have not been thoroughly investigated in photosynthetic systems, including cyanobacteria. The highly disordered protein Alr0806 from *Anabaena* shows partial homology to the HLIP from *Synechococcus* sp. JA-3-3Ab (**10, Suppl Fig 1C**), but surprisingly does not share significant homology to other HLIPs from *Anabaena*. Among the filamentous cyanobacteria, the absence of Alr0806 in primitive cyanobacteria (e.g., Thermoleptolyngbia or *Pseudanabaena*), but not in more recent species such as *Anabaena* indicates it arose later during cyanobacterial evolution. Although, the presence of the Alr0806 homologs broadly aligned with major phylogenetic clades, several close taxa showed divergence, suggesting that Alr0806 underwent lineage-specific retention, possibly to assist stress adaptation and cellular complexity in more advanced species.

The N-terminal region of Alr0806 protein, identified in the cell surface fraction of *Anabaena* PCC 7120, was determined to be TNEPVKRATDSSEXA (**Yoshimura et al., 2012)**. This sequence begins precisely 22 amino acids downstream from the database-annotated start site of Alr0806 (**Figure 1**). In their study, Yoshimura et al., proposed the presence of a signal peptide (IFKKHNLVLEVL TDFLTRTDFM/T), whose cleavage resulted in generation of the smaller Alr0806 protein with the above-mentioned N-terminal sequence. However, our study confirmed that the *alr0806* transcript originates downstream of the translational start annotated in databases (**Figure. 1C)**. Moreover, MSA clearly revealed the sequence similarity of Alr0806 and its cyanobacterial/bacterial homologs to commence from the methionine located 22 amino acids away from the annotated translational start (**Suppl Fig 1B**). These evidences support our finding that the ATG codon downstream of the annotated ‘GTG’ start codon is the actual start codon and the use of this ATG results in synthesis of 135 residue Alr0806 protein in *Anabaena* (**Figure 1**).

The abnormal movement of the Alr0806 protein on SDS PAGE (**Rai et al., 2013, Yoshimura et al. 2012**), was attributed to its association with entities such as lipids and sugars or formation of tight dimers (**Yoshimura et al., 2012)**. However, it is now established that IDPs, which lack well-defined structure, do not conform to the typical rod-like shape that SDS-PAGE assumes for globular proteins, and this structural flexibility usually leads to anomalous migration patterns. For example, the highly acidic IDP Gir2 from *Saccharomyces* also shows anomalous slow migration due to its high content of negatively charged residues (**Alves and Castilho, 2005**). Thus, the higher apparent mol wt. of Alr0806 on SDS PAGE is due to its high negative charge contributed by the long stretches of disordered regions in the protein (**Figure 2**). Highly acidic proteins tend to repel SDS molecules, leading to insufficient binding of SDS, which results in decreased electrophoretic mobility (**Figure 2**).

IDPs may undergo significant conformational changes near their pI due to increased hydrophobic interactions and transition from a disordered to a more ordered state because of reduction in net charge, which minimizes repulsive forces (**Tedeschi et al., 2017**). At its pI (pH 4.0) or on treatment with helix stabilizing agent (TFE), the conformation of Alr0806 altered, reflecting enhanced helical content (**Figure 2**). Generally, the multiple conformations exhibited by IDPs enables them to engage in a variety of interactions with other biomolecules (**Lindstrom et al., 2018**) often with high specificity but low affinity (**Olsen et al., 2017**). Thus, IDPs can effectively function as an interaction “Hub” (highly connected nodes) in relation to a specific conformational ensemble (**Yruela and Neira, 2020**). As shown in **Figure 3**, Alr0806 formed an extensive protein-protein-interaction network with over 50 proteins involved in major metabolic pathways.

Alr0806 likely leverages its capability to undergo disorder-to-order transitions (**Figure 2**) to perform critical biological functions. Some IDPs such as dehydrins and LEA play an important role in plant stress responses, particularly through their chaperone functions (**Tompa and Kovacs, 2010, Kovacs et al., 2008**). Alr0806 also prevented aggregation of other proteins at high temperatures as well as under conditions of dehydration, underscoring its ability to function as a chaperone (**Figure 3**). Heavy metals drought and chemicals like butachlor not only denature cellular proteins but also they also trigger oxidative stress, inflicting damage to vital biomolecules such as DNA. Studies have shown *Anabaena* responds to these stresses by ramping up the production of chaperones (GroEL/PPIase), DNAase (Alr3199) and ROS scavenging proteins such as the Mn-catalase, Alr3090 (**Chakravarty et al., 2016**, **Agrawal et al., 2014**). Interestingly, all the above-mentioned proteins (i.e. Alr0806, Alr3090, & Alr3199) are strongly induced in response to salinity (**Chakravarty et al., 2016**, **Rai et al.,2013**). This structural response suggests a coordinated activation of the cellular machinery necessary to combat the multifaceted challenges of protein unfolding, DNA damage, dehydration, and oxidative stress. Interestingly, genes encoding these proteins repressed by the ferric uptake regulator A (FurA) in *Anabaena*, implying that stress responsive proteins which regularly intersect to overcome environmental perturbations, may be co-ordinately regulated. Moreover, the heat shocked *Anabaena* filaments show improved tolerance to salt stress (**Srivastava et al, 2023**), indicating that elevated production of chaperones during heat stress reduces the debilitating effects of salinity. Thus, the chaperone activity of the salt-induced Alr0806 is likely to protect intracellular proteins from unfolding or aggregation caused by salinity.

To summarize, the current study shows Alr0806 to be an extremely thermostable, multiple stress-induced, intrinsically disordered protein that is capable of disorder-to-order transitions. The dynamic nature of Alr0806 enables it to function as a chaperone by facilitating proper folding and preventing aggregation of target proteins. The structural flexibility of Alr0806 also enables it to function as a hub protein, which interacts with more than 50 cellular proteins. Notably, Alr0806 is not only required by *Anabaena* for optimum growth under standard conditions, but also for augmenting stress resilience under adverse conditions. Elucidation of Alr0806 function has illuminated the physiological role of this class of proteins and opened newer avenues for biotechnological applications aimed at improving stress tolerance in photosynthetic systems.

## Author contribution

MB and AB: conceptualization; DTH: investigation; data curation, methodology; DTH, AB and MB: formal analysis; DTH and MB: writing - original draft; AB: reviewing and writing final draft; MB and AB: supervision

## Acknowledgement

We thank Dr. Hema Rajaram, Head MBD for her support during the course of this work.

## Data Availability Statement

All supporting data are included within the main article and its supplementary files.

## Materials and Methods

### Organism and growth conditions

BG-11 liquid medium, with combined nitrogen (17 mM NaNO3), pH 7.2 (Castenholz, 1988) was used to grow axenic cultures of Anabaena PCC7120 strains under continuous illumination (30 µE m−2 s−1), with or without shaking (100 r.p.m.), at 27°C ± 2°C. Content of chlorophyll a ml−1 was estimated as described by **Mackinney (1941**) and served as a measure of growth. *Escherichia coli* cells were grown in Luria–Bertani (LB) medium in the presence of appropriate antibiotics at 37°C with shaking at 150 r.p.m. The antibiotic neomycin (Nm) was used at 10 µg ml−1 (Nm10) and 25 µg ml−1 (Nm25) in BG-11 liquid media and BG-11 agar plates respectively for the recombinant *Anabaena* PCC7120. Antibiotics used for E. coli were 34 µg chloramphenicol ml^−1^ (Cm34), or 100 µg carbenicillin ml^−1^ (Cb100).

### Evolutionary analysis

The 16S ribosomal RNA (rRNA) gene sequences for 56 representative cyanobacteria encompassing diverse orders were retrieved from either the Integrated Microbial Genomes & Microbiomes (IMG) database (https://img.jgi.doe.gov/) or the National Center for Biotechnology Information (NCBI) GenBank (https://www.ncbi.nlm.nih.gov/). Multiple sequence alignments (MSAs) were generated using MAFFT v7.508 (**Katoh et al., 2019**) with the L-INS-i algorithm for accurate alignment of homologous regions. Subsequently, a maximum likelihood (ML) phylogenetic tree was constructed using IQ-TREE v1.6.12 (**Trifinopoulos et al., 2016**). ModelFinder (**Kalyaanamoorthy et al., 2017**) was employed to identify the most appropriate substitution model (GTR+F+I+G4) for tree reconstruction, and robustness was assessed with 1000 ultrafast bootstrap replicates. Putative orthologs for Alr0806 proteins were identified using a homology search approach. Protein sequence of Alr0806 was retrieved from the UniProt (https://www.uniprot.org/). Subsequently, a BLASTP search against the non-redundant protein sequence database (nr) was performed using NCBI’s BLAST+ suite **(Camacho et al., 2008)** with the following parameters: E-value threshold of 1×10^-6 (-evalue 1e-6), a maximum target sequence of 5000 (-max_target_seqs 5000), and a minimum identity filter of 25% to ensure a significant degree of sequence similarity (**Ward et al., 2014**).

### RNA isolation and analysis

Isolation, electrophoresis of RNA and Northern blotting analysis was performed as described earlier (Ballal and Apte, 2005). For RT-PCR analyses, RNA was quantified on a Nano Drop spectrophotometer (Thermo Fisher Scientific) and subjected to cDNA synthesis (**Banerjee et al., 2021**). To obtain, DNA-free RNA, ∼1 µg RNA was treated with 1 U of RNase-free DNase (Sigma Aldrich). Subsequently 500 ng RNA was subjected to reverse transcription using PrimeScript™ 1^st^ strand cDNA Synthesis Kit (Takara Bio Inc.). KAPA SYBR^®^ FAST qPCR Master Mix (2X) Kit (KAPABIOSYSTEM) was used for qPCR reaction on Master cycler ep realplex Real-time PCR instrument (Eppendorf, Germany). Thermal cycling conditions were standardized and fixed as 94◦ C (2 min), 45 cycles of 94◦ C (10 s), 60◦ C (15 s), 68◦ C (20 s). 100 ng of total RNA was used for each reaction in triplicate (with amplicon size <250 bp). Primers for the *alr0806* gene were designed using the primer 3 software. PCR amplification was confirmed by the melting curve analysis or by resolving the products on 2.0 % agarose gels. For each gene (-RT) control was kept for every reaction. C_t_ values, which represents the absolute quantification of the expression was further used to calculate the fold change. RNaseP RNA gene (*rnpB*), which is a housekeeping gene in *Anabaena* PCC 7120, was employed as an internal control (**Banerjee et al., 2021**). The relative-fold change quantification of a target gene in 100 mM NaCl-treated cells in comparison to the control cells is expressed in terms of 2^−ΔΔCt^ values after normalization with the internal control (*rnpB*). Experiments were repeated twice.

### Construction of plasmid for knocking down *alr0806* and its conjugation into *Anabaena* PCC 7120

A CRISPR–dCas9-based plasmid for knocking down *alr0806* was constructed following the strategy described by **Kalwani et al. 2022**. A sgRNA specific to *alr0806* was inserted into the XmaI–SalI site of the CRISPR–dCas9 based knocking down vector (pKD4641) to selectively target *alr0806* transcription (construct designated as pKD0806). A plasmid lacking the specific sgRNA (designated pKDScr) was employed as the control plasmid generated. These constructs were introduced into *Anabaena* sp. PCC 7120 by triparental conjugation as described by **Kalwani** et al. (2022).

### Disorder prediction and 3D structure of Alr0806

PONDR server was employed to predict the intrinsic disorder of Alr0806. The amino acid sequence of Alr0806 retrieved from the KEGG database was used as input for the analysis. Three-dimensional structures of both the Alr0806 protein was predicted using the AlphaFold server. The liquid–liquid phase separation (LLPS) propensity of Alr0806 was predicted by the FuzDrop server.

### Cloning, Expression and purification of Alr0806

The *alr0806* ORF was PCR-amplified with forward primer and reverse primer. employing *Anabaena* PCC 7120 genomic DNA as a template. Restriction-enzyme sites for Nde1 and BamHI were incorporated in the forward and the reverse primer, respectively. The reverse primer also had six His codons (shown in bold) followed by a stop codon. The amplified products were purified, digested with the corresponding restriction enzymes, and ligated into a pET21a expression vector under the control of the T7 promoter. DNA sequencing was performed to ensure the nucleotide sequence integrity of *alr0806*. For production of the tag-less Alr0806 protein, the *alr0806* ORF was PCR-amplified using the primer pair F & R (lacking the His-tag sequence. The amplified DNA was inserted into pET21a as described above.

For protein expression, verified plasmids encoding either His-tagged or tag-less Alr0806 were transformed into *E. coli* BL21 (DE3) pLysS cells. Transformed cells were cultured in Luria–Bertani (LB) medium supplemented with appropriate antibiotics at 37 °C with shaking at 180–200 rpm. When the cultures reached OD_600_ of approximately 0.5–0.6, expression of the recombinant protein was induced with 1 mM isopropyl β-D-1-thiogalactopyranoside (IPTG). After incubation at 20 °C (overnight) cells were harvested by centrifugation at 4 °C and stored at −70 °C until further processing. Cell pellets were resuspended in lysis buffer [20–50 mM Tris-HCl (pH 8.0), 200 mM NaCl, and 5 mM imidazole], lysed by sonication on ice and clarified by centrifugation at 15,000 rpm for 30 min at 4 °C.

For purification of the His-tagged Alr0806, the clarified supernatant was incubated with Ni²–NTA agarose resin pre-equilibrated with lysis buffer. After binding, the resin was washed with buffer containing 10–30 mM imidazole to remove non-specifically bound proteins. The target protein was eluted using 100–250 mM imidazole. Eluted fractions were analyzed by SDS–polyacrylamide gel electrophoresis (SDS–PAGE), and fractions containing purified protein were pooled and dialyzed overnight against 20 mM Tris-HCl (pH 8.0) to remove imidazole.

For purification of the tag-less Alr0806, the clarified lysate was subjected to heat treatment at 100oC (30 min) followed centrifugation (15,000 g, 30 min at 4 °C) to remove the heat denatured proteins and the supernatant was subjected to anion-exchange chromatography. The protein solution was loaded onto a column pre-equilibrated with 20 mM Tris-HCl (pH 8.0). After thorough washing, the bound protein was eluted using a linear gradient of NaCl (0–1 M). Fractions were analyzed by SDS–PAGE, pooled on the basis of purity, and dialyzed overnight against 20 mM Tris-HCl (pH 8.0) to remove NaCl.

### SDS–PAGE Analysis

Protein samples collected at various purification stages were analyzed by resolving them on SDS– (12%) polyacrylamide gels followed by staining with. Molecular weight estimation was performed using protein standards.

### Dynamic Light Scattering (DLS)

Dynamic light scattering measurements were performed to determine the hydrodynamic radius of the purified Alr0806. Experiments were conducted at 25 °C using a Zetasizer ZS90 (Malvern Instruments Ltd.). Protein samples prepared in 20 mM Tris-HCl (pH 8.0) were analysed in standard 1ml quartz cuvettes provided with the instrument. Measurements consisted of multiple acquisitions, and the average hydrodynamic diameter was calculated using instrument software.

### Circular Dichroism (CD) Spectroscopy

Far-UV circular dichroism spectra were recorded to evaluate the secondary structural content of Alr0806. Measurements were performed at 25 °C in the wavelength range of 195–260 nm using a 1 mm path-length quartz cuvette. Protein samples at concentrations of 0.1–0.3 mg/mL were prepared in 20 mM Tris-HCl buffer. Each spectrum represented the average of multiple scans and was corrected for buffer baseline. Observed ellipticity was expressed in millidegrees (mdeg), and secondary structure content was estimated using standard deconvolution software.

### Intrinsic tyrosine Fluorescence Spectroscopy

Intrinsic tyrosine fluorescence measurements were carried out to assess tertiary structure and conformational stability. Protein samples were excited at 275 nm, and emission spectra were recorded between 290 and 400 nm at 25 °C. Changes in emission maxima and fluorescence intensity were monitored under native and different pH conditions.

### Limited Proteolysis

Limited proteolysis experiments were performed to examine the structural flexibility of Alr0806. Purified protein was incubated with sequencing-grade trypsin at a different enzyme-to-substrate ratio such as 1:1000, 1:5000 and 1:10000 (w/w) in 20 mM Tris-HCl (pH 7.5–8.0) at 25 °C. Aliquots were withdrawn at defined time intervals (2, 5, 10, 15, 30, 60, 120 & 240 min) and the reaction was terminated by the addition of SDS loading buffer followed by heating at 95 °C for 5 min. Proteolytic fragments were resolved on 12–15% SDS–PAGE and visualized by staining. The digestion pattern was analysed to assess protease susceptibility and structural accessibility.

### Chaperone Activity Assay

The chaperone-like activity of Alr0806 was evaluated by monitoring its ability to suppress thermal aggregation of malate dehydrogenase (MDH). Aggregation assays were carried out in 20 mM Tris-HCl (pH 7.5) at elevated temperature (42–45 °C). MDH (approximately 0.2–0.3 mg/mL) was incubated either alone or in the presence of increasing molar ratios of Alr0806 (1:1, 1:2, 1:5 & 1:10). Protein aggregation was monitored by measuring light scattering at 360 nm using a spectrophotometer. The increase in absorbance over time reflected MDH aggregation. Suppression of aggregation in the presence of Alr0806 was calculated relative to MDH alone, which served as a control. All experiments were performed in triplicate.

### Dehydration Protection Assay

The dehydration protection capacity of Alr0806 was assessed by determining its ability to preserve MDH enzymatic activity following desiccation stress. MDH was incubated alone or with ALR0806 at various molar ratios (1:1, 1:2, 1:5 & 1:10) in 20 mM Tris-HCl (pH 7.5). Samples were subjected to dehydration by vacuum desiccator at room temperature. Subsequently, samples were rehydrated with the original buffer volume. Residual MDH activity was determined spectrophotometrically by monitoring the oxidation of NADH at 340 nm in the presence of oxaloacetate. Enzyme activity after dehydration was expressed as a percentage relative to non-dehydrated controls. The protective efficiency of ALR0806 was calculated by comparing retained MDH activity in the presence and absence of ALR0806.

### Protein-protein interaction network and pathway analysis

PPI data for Alr0806 was retrieved from the cyanomapDB database (Peng et al., 2023). Cytoscape software (https://cytoscape.org) (Shannon et al., 2003) was utilized to visualize and analyze the interaction networks based on the extracted PPI data. For pathway analysis, the genes involved in PPI network were uploaded to ShinyGO 0.80 (http://bioinformatics.sdstate.edu/) for Gene Ontology (GO) enrichment analysis.

### PAM Fluorimeter

Chlorophyll fluorescence was measured using the Dual-PAM-100 fluorimeter (Heinz Walz GmbH). Measurements (measuring light: 470 nm) were performed using the automated programme (Dual PAM v3.20) and PSII activities were quantified by monitoring changes in the chlorophyll *a* fluorescence. Before F_V_/F_M_ measurements, the samples were dark adapted for 15 min. The relative electron transport rates (rETRs) of PSII [rETR(II)] were recorded during the measurement of the light response curve with increasing illumination. In light response curve, sequence of illumination steps was carried out with increasing saturation pulse of photosynthetically active radiation (PAR). Experiments were repeated at least two times.

### Growth analysis

Growth of control (An-dCas9) or the *alr0806* knockdown (An-KD0806) *Anabaena* cells with or with 100 mM NaCl treatment was measured by monitoring the amount of chlorophyll a at the depicted time points. For estimation of the chlorophyll a content, *Anabaena* cells present in 1 ml culture were harvested by centrifugation, 1 ml of 90% methanol was added to the cell pellet, the mixture was thoroughly vortexed to extract the chlorophyll a and A_665_ was monitored with a spectrophotometer. All the experiments were repeated at least three times.

## Figure legends

Supp Figure 1: (A) quantitative reverse transcriptase PCR to monitor levels of the *alr0806* transcript under NaCl or NH_4_Cl. (B) Multiple sequence alignment of the cyanobacterial Alr0806 homologs.

Supp Figure 2: (A) Liquid–liquid phase separation (LLPS) propensity of Alr0806 predicted by the Fuzdrop server. (B) Diagram-of-state representation of Alr0806 based on the distribution of charged amino acids within its sequence. (C) Purification of His-tagged Alr0806 using Ni–NTA affinity chromatography. Proteins were eluted with increasing imidazole concentrations (100–250 mM). (D) CD spectra of Alr0806 recorded at different temperatures. (E) Boiled cell lysate containing the Alr0806 protein was passed through an anion-exchange column and eluted using an increasing NaCl gradient. Different elution fractions (F1–F8) were analyzed on Coomassie-stained SDS-Polyacrylamide gels. M denotes the molecular weight marker. Inset showing the MALDI–TOF mass spectrometric profiles of fractions F5 and F6. (F) Native PAGE analysis of purified Alr0806 protein. (G) DLS analysis displays a single, narrow peak corresponding to a hydrodynamic radius of 7nm. (H) Intrinsic tyrosine fluorescence emission spectrum of Alr0806 protein under different pH condition Supplementary Figure 3: (A) Pathway enrichment analysis of Alr0806-interacting proteins using ShinyGO 0.8 (http://bioinformatics.sdstate.edu/). (B) PPI network of Alr0806 constructed using interaction data from Xu et al. (2022). Orange node represents Alr0806, while grey nodes represent interacting proteins.

